# An Interpretable Framework for Clustering Single-Cell RNA-Seq Datasets

**DOI:** 10.1101/191254

**Authors:** Jesse M. Zhang, Jue Fan, H. Christina Fan, David Rosenfeld, David N. Tse

## Abstract

**Background:** With the recent proliferation of single-cell RNA-Seq experiments, several methods have been developed for unsupervised analysis of the resulting datasets. These methods often rely on unintuitive hyperparameters and do not explicitly address the subjectivity associated with clustering.

**Results:** In this work, we present DendroSplit, an interpretable framework for analyzing single-cell RNA-Seq datasets that addresses both the clustering interpretability and clustering subjectivity issues. DendroSplit offers a novel perspective on the single-cell RNA-Seq clustering problem motivated by the definition of “cell type,” allowing us to cluster using feature selection to uncover multiple levels of biologically meaningful populations in the data. We analyze several landmark single-cell datasets, demonstrating both the method’s efficacy and computational efficiency.

**Conclusion:** DendroSplit offers a clustering framework that is comparable to existing methods in terms of accuracy and speed but is novel in its emphasis on interpretabilty. We provide the full DendroSplit software package at https://github.com/jessemzhang/dendrosplit.

## 1. Background

In recent years, single-cell RNA-Seq has proven to be a powerful approach for studying biological samples in various settings^1^. Scientists have leveraged this technology to shed light on how cells differentiate^2–6^, investigate known cell types^7–10^, and discover new cell types and gene patterns^11–17^. These efforts have yielded a plethora of diverse datasets sharing characteristics such as missing entries (drop-out events) and high dimensionality. Additionally, technological breakthroughs such as droplet encapsulation, molecular barcoding, and cheap parallelization have produced datasets involving tens of thousands and even millions of cells^17–22^. After obtaining such datasets, scientists are often interested in clustering the high-dimensional points corresponding to individual cells, ideally recovering known cell populations while discovering new and perhaps rare cell types. While the definition of a cell type is not precise^23^, biologists agree that gene expression levels are highly relevant. With gene expression dictating protein expression (and hence cellular function), identifying the genes that distinguish a cell type is of paramount importance. Therefore from a computational perspective, there are two key problems in downstream analysis: 1) clustering and 2) feature selection, also known as differential expression.

General-purpose clustering algorithms such as *K*-means, DBSCAN^24^, affinity propagation^25^, and spectral clustering^26^ have performed well for several single-cell datasets^27^. In order to achieve good performance, however, the datasets often need to be carefully preprocessed, and the algorithms require non-intuitive hyperparameter tuning. For example, both K-means and spectral clustering require choosing the desired number of clusters, DBSCAN requires choosing the max distance between two samples in the same neighborhood, and affinity propagation requires choosing both a preference parameter for determining which points are exemplars and a damping parameter for avoiding numerical oscillations. To address specific computational challenges of single-cell RNA-Seq datasets, researchers have developed a wide array of application-specific clustering algorithms^28–34^ and packages for end-to-end analysis^21,35–39^. Regardless of which set of these tools one uses, finding the right approach for clustering a specific dataset requires careful design of the computational workflow, but often finding a good combination of clustering algorithm and hyperparameters is time-consuming and difficult. Additionally, none of these approaches explicitly addresses the inherent subjectiveness behind clustering, which stems from the potential existence of subtypes and sub-subtypes.

With an emphasis on interpretability and ease of exploratory analysis, we introduce DendroSplit, a framework for clustering single-cell RNA-Seq data. In addition to speed, the framework has the following advantages:

- Gene-based justification for all decisions made when generating clusters
- Interpretable hyperparameters
- Ability to cheaply produce multiple clusterings for the same dataset
- Ease of incorporation into existing single-cell RNA-Seq workflows

At a high level, the approach leverages a feature selection algorithm to generate biologically meaningful clusters. The end-to-end DendroSplit workflow is illustrated in Figure 1A. After preprocessing the *N* × *M* expression matrix *X* (where *N* and *M* represent the number of cells and genes, respectively), we generate the *N* × *N* distance matrix *D*. We use hierarchical clustering to iteratively group cells based on their pairwise distances, obtaining a dendrogram, a tree-like data structure illustrating how grouping was performed. The split step starts at the root of the tree. Each node in the dendrogram represents a potential partitioning of a larger cluster into two smaller ones. If this “split” results in two adequately separated clusters (according to a metric we call the **separation score**), the split is deemed valid and the algorithm continues on the new clusters. Otherwise, the algorithm terminates for the subtree below the node. After the split step, DendroSplit performs a pairwise comparison of the resulting clusters, repeatedly merging clusters until all clusters are sufficiently separated. The merge step counteracts the greedy nature of hierarchical clustering, allowing DendroSplit to compare clusters that may have incorrectly ended up far away from one another in the dendrogram. The overall approach involves two intuitive hyperparameters: the separation score threshold for accepting a split, and the separation score threshold for accepting a merge.

**Figure 1.**
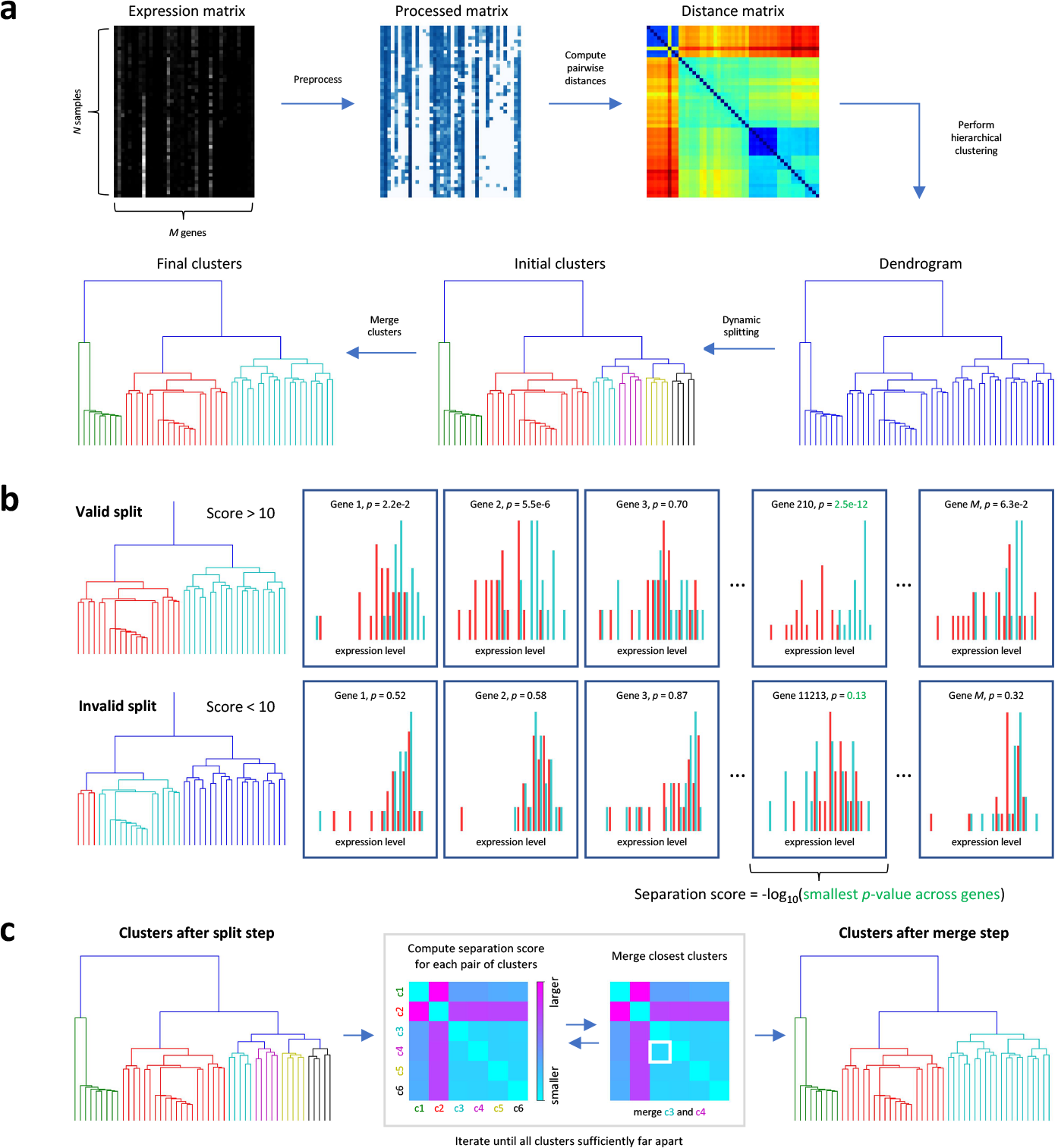
Overview of DendroSplit. **(a)** The workflow starts by preprocessing a *N* × *M* matrix of gene expressions before computing cell-cell pairwise distances, resulting in a *N* × *N* distance matrix. The distance matrix is fed into a hierarchical clustering algorithm to generate a dendrogram. The dynamic splitting step involves recursively splitting the tree into smaller subtrees corresponding to potential clusters. Finally, the subtrees are merged together during a cleanup step to produce final clusters, **(b)** A split corresponds to the partitioning of a larger cluster into two smaller clusters. A split is only deemed valid if the separation score, a metric for how well-separated two populations are, is above a predefined split threshold. Leveraging biological intuition, we rank how well each gene distinguishes the two subpopulations based on independent Welch’s *t*-tests. We use the — log of the smallest *p*-value obtained as our separation score due to its interpretability and practical effectiveness. A split threshold of 10 would work for the example shown here, **(c)** During the merge step, the clusters obtained from the split step are compared to one another using pairwise separation scores. If the closest two clusters are not sufficiently far apart based on a predefined merge threshold, they are merged together and the process is repeated. When all clusters are sufficiently far apart, the algorithm terminates and the final labels are output.

We use the term “framework” to underline how specific design choices for certain components in the workflow such as the separation score will result in different clustering “methods.” Our choice of separation score is motivated by a key assumption: **if two cell populations are of different types, then there should exist at least one gene that is differentially expressed between the two populations**. Given a candidate split in the dendrogram, we perform a Welch’s *t*-test for each gene. The separation score is – log(*p*_min_) where *p*_min_ represents the smallest *p*-value achieved (Figure 1B), and we will be using this definition of separation score for all experiments presented in this work. We demonstrate that the deterministic method outlined in Figure 1 is applicable to a wide variety of single-cell datasets. We show how DendroSplit can help us investigate the most significant genes considered at each split or merge, providing insight for how clusters are generated. Finally, we show how DendroSplit can cheaply generate several clusterings for different hyperparameter values.

Some clustering approaches similar to DendroSplit exist in literature. For example, the most common method of generating clusters from a dendrogram involves simply cutting the dendrogram horizontally at some fixed height. This rigid approach often fails to generate meaningful clusters for more complex datasets. The Dynamic Tree Cut algorithm^40^ adds significant flexibility and processes the dendrogram based on an adaptive cut. Though it does not explicitly use a dendrogram, the backSPIN algorithm^12^ also uses cell-cell similarities to perform iterative splitting. Unlike DendroSplit, both of these algorithms require choosing unintuitive hyperparameter cutoffs based on nuanced criteria. The most similar clustering approach was used by Lake et al.^15^ for analyzing their human brain single-cell dataset. Their approach fits into the DendroSplit framework, using a separation score based on random forests. This separation score, compared to the separation score mentioned above, has an element of randomness, is significantly more computationally expensive, and requires less intuitive hyperparameter choices.

## Implementation

### Distance metric

For all single-cell datasets, we used the correlation distance. The correlation distance between **x**_*i*_ and **x**_*j*_ corresponding to cells *i* and *j* is

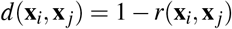

where *r* is the Pearson correlation coefficient. Therefore d is bounded between 0 and 2. This distance metric has the advantage of being agnostic to both shift and scale, making it robust to certain biases we would expect to vary across datasets. As a caveat, the distance metric has the disadvantage of depending on the number of zeros, and therefore the distance between two cells before and after gene filtering may be different due to removal of entries equal to 0. For all experiments in this work, distance matrix computations were parallelized on 32 cores and computed using the `scikit-learn` Python package^41^.

### Hierarchical clustering

DendroSplit performs hierarchical clustering using the Scipy Python package^42^. One source of ambiguity for hierarchical clustering lies in the method for determining the distance between two clusters. We found that the “complete” method produces the best results, and this is the method used for all experiments reported below. For this method, the distance between two clusters is equal to the largest distance between a point from the first cluster and another point from the second cluster.

### Separation score

The separation score effectively serves as a distance metric between two clusters, quantifying how different they are (see the supplementary material for further discussion). The cell-type assumption discussed in the Background section can also be phrased as: if two cell populations are of different types, then projection along one of the *M* axes should result in two distinguishable point clouds. For all experiments performed in this work, we defined the separation score between the *N*_1_ × *M* population **X** and the *N*_2_ × *M* population **Y** as

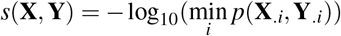

where *p*(**X**_.*i*_,**Y**_.*i*_) represents the *p*-value achieved using a Welch’s *t*-test for gene *i*. **X**_.*i*_ represents the ith column of **X** corresponding to the expression of gene *i* in population **X**. Welch’s *t*-test is similar to Student’s *t*-test but is more reliable when the two populations have unequal variance and size^43^. Compared to other differential expression approaches, Welch’s *t*-test is computationally cheap.

As an implementation note, if for a given split two or more genes have the exact same score, these genes are ranked by the magnitude of the *t* statistic. We note that because we are using Welch’s *t*-test rather than Student’s *t*-test, the degrees of freedom associated with each test is different, and hence outputting the largest *t* statistic is only approximately sound. Two genes may have the exact same score for larger datasets and for splits near the root of the dendrogram where *p*-values may be quite small, resulting in an underflow issue and a score of ∞.

### Handling singletons

In addition to the split and merge thresholds, the two major hyperparameters discussed in the Background section, the DendroSplit framework can also be customized using three minor hyperparameters. These three hyperparameters are relevant for finding singletons (clusters containing one point), which are analogous to outliers.

Two of these hyperparameters are relevant for the split step. The first is the minimum cluster size. During a split, if one of the two candidate clusters contains less points than the minimum cluster size, that cluster is disbanded (each of its points are labeled as “Singleton”) and the algorithm continues on the other candidate. The second hyperparameter is the disband percentile. If a candidate split does not produce a subtree that meets the minimum cluster size requirement or if the candidate split does not achieve a high enough separation score, we look at the pairwise distances amongst samples in this final cluster. If all of them are greater than a certain percentile of distances in *D*, the original *N* × *N* distance matrix, then all points in this final cluster are marked as singletons. For all experiments performed in this work, the minimum cluster size was set to 2 (the smallest value) and the disband percentile was set to 50.

Before merging clusters, each singleton obtained during the split step is assigned to the same cluster as its nearest neighbor. If the distance between a singleton and its nearest neighbor is greater than a certain percentile of all pairwise distances in *D*, then the singleton remains unclassified. This percentile is the third minor hyperparameter and was set to 90 for all experiments performed in this work.

### Hyperparameter Sweeping

When choosing hyperparameters for DendroSplit, a relatively small split threshold such as 20 and a merge threshold set to half the split threshold often yields reasonable initial results. The DendroSplit approach can also rapidly generate several clusterings based on different split thresholds. Since DendroSplit saves the *p*-values and cell IDs considered at each split, we can obtain several split-step clustering results by exploiting the fact that the clusters generated with a smaller score threshold partition the clusters generated with a larger score threshold. The merge threshold can then be chosen by looking at pairwise separation scores between clusters.

## Results

### Data preprocessing

For all single-cell datasets, we apply a logarithmic transformation log_10_(*X* + 1) to the raw expression levels. We analyze 9 datasets in this paper. For each dataset, genes that have 0 expression across all cells were removed. Additionally, all datasets consisting of over 1000 cells undergo feature selection based on the method proposed by Macosko et al.^20^ The *M* genes are sorted into equal-sized bins depending on their mean expression values. Within a bin, genes are *z*-normalized based on their dispersions, where the dispersion for a gene is defined as the variance divided by the mean. Only genes corresponding to a *z*-score above a certain cutoff are retained. For the Zeisel et al.^12^, Birey et al.^17^, and Zheng et al.^21^ datasets, we use DendroSplit’s default setting of 5 bins with a *z*-cutoff of 1.5. For the Macosko et al. dataset, we first remove cells with less than 900 counts across all genes just like in Wang et al.’s^30^ approach, reducing the original 44808 cells to 11040. We then use Macosko et al.’s gene-filtering settings of 20 bins with a *z*-cutoff of 1.7. For the Zheng et al. dataset, reducing the number of genes results in several of the original 68579 cells having few and even 0 counts across all genes. We remove cells with less than 50 counts across all genes, resulting in 17426 cells, and again filtered out genes with 0 counts across all remaining cells. We also experimented with standardizing all log-transformed genes to have 0 mean and unit variance across all cells, but the increased computational overhead did not yield better results.

### Adjusted Rand index

The adjusted Rand index (ARI) is used to quantify how our clustering results match another given set of labels. The ARI ranges from 0 for poor matching to 1 for perfectly matched labels. For a set of *n* elements, we let *X* and *Y* represent two partitions of the elements. *X_i_* represents the set of elements in partition *i* according to *X*. The adjusted Rand index is defined as:

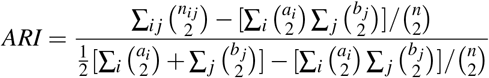

where *n_ij_* is the number of elements in common between *X_i_* and *Y_j_*, *a_i_* = ∑*_k_n_ik_*, and *b_j_* = ∑*_k_n_kj_*.

### Ground-truth datasets

To test the effectiveness of the DendroSplit framework, we first test the approach on datasets where the ground truth is known.

#### Synthetic datasets

Figure 2 shows the performance of DendroSplit on four synthetic datasets^44^. Since the 2-dimensional data points have clear, intuitive clustering structure, pairwise Euclidean distance is a natural choice. Figure 2A shows the exploratory power of DendroSplit on a toy dataset of oddly-oriented clusters. Because DendroSplit saves the information gathered at each valid split, we can easily investigate how the clustering was performed. At a given split, we can identify the points that went into each partition and look at the partition-specific distributions of the feature that validated the split. Thus the true advantage of DendroSplit is in its ability to justify its behavior with interpretable results. Figure 2B shows that DendroSplit has the power to uncover several clusters especially when the distance metric (Euclidean) suits the type of data (2-D Gaussian balls). Figure 2C emphasizes that, like other methods, DendroSplit cannot automatically overcome poor choices in preprocessing and distance metric selection.

**Figure 2.**
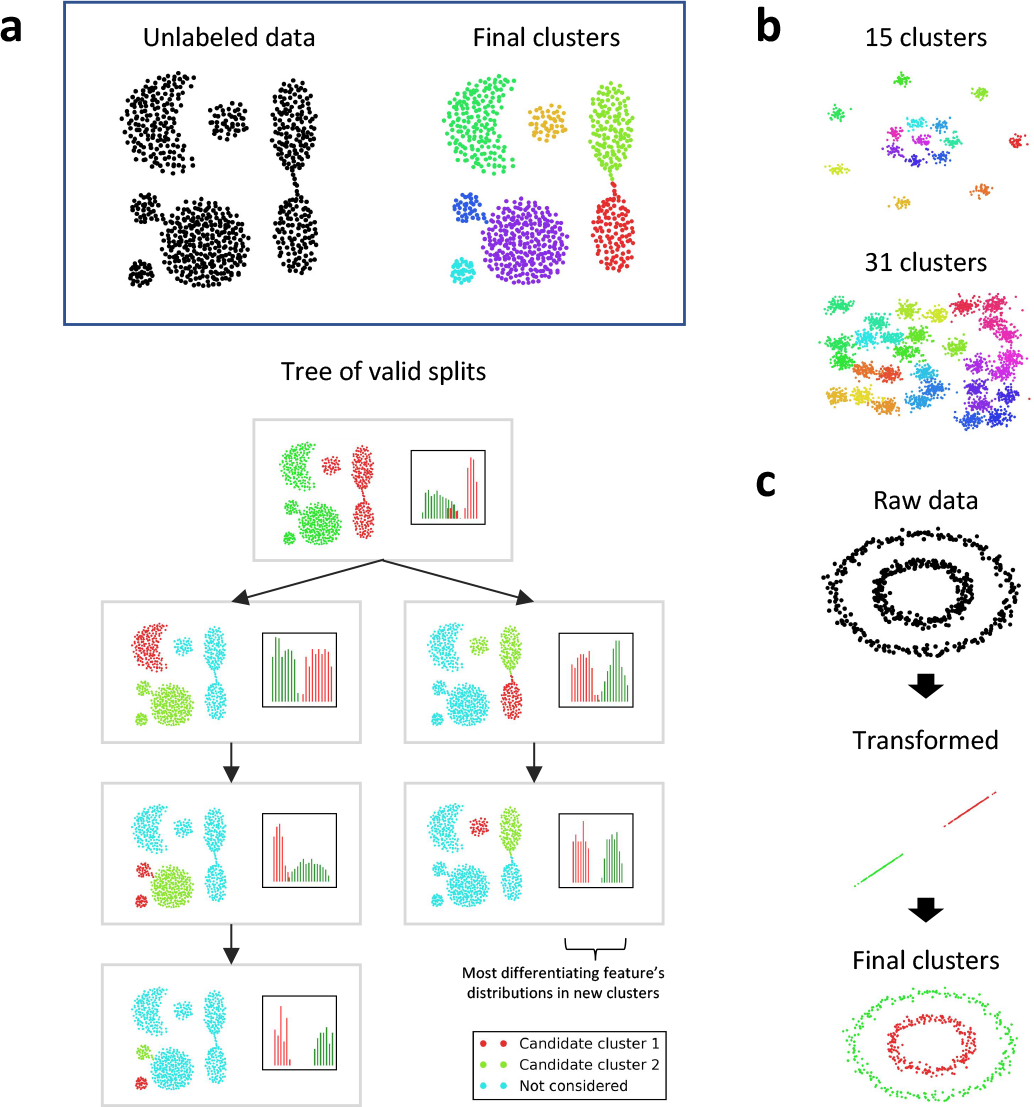
Synthetic datasets. **(a)** The DendroSplit approach is applied to a synthetic 2-dimensional dataset where pairwise distances are equal to the Euclidean distances between points. The dendrogram splitting process can be visualized using a tree, and each box in the tree represents a step in the algorithm where a larger cluster is partitioned into the red and green clusters. For each of the two features (dimensions), the split is evaluated based on the distributions of that feature within the candidate clusters. Teal points are “background” points and not considered for a given step, **(b)** DendroSplit is evaluated on two other 2-dimensional synthetic datasets and recovers the correct number of clusters both times. Euclidean distance is used, **(c)** We note that DendroSplit cannot overcome poor preprocessing and distance metric selection. Directly computing Euclidean distances for the points in the concentric circle dataset would yield poor performance, but using Euclidean distance after some preprocessing (e.g. mapping each point to its distance from the center) yields the correct results. For the examples shown here, the merge thresholds are 10, and the split thresholds are 40 for **(a)** and 30 for **(b,c).**

#### Single-cell RNA-Seq datasets

Figure 3 shows the performance of DendroSplit on four single-cell RNA-Seq datasets featuring high-quality labels. Kiselev et al.^34^ refers to these datasets as “gold standards.” We chose four datasets with varying amounts of cells, genes, and total clusters to understand how they affect the behavior of DendroSplit. We see that when *N* is on the order of 100s, the runtime is widely determined by *M*, the number of independent Welch’s *t*-tests that must be performed at every split. Figure 3 shows that for the Biase et al.^2^, Pollen et al.^8^, and Kolodziejcyk et al.^5^ datasets, most of the final ARI is achieved after the split step. Therefore most of the information captured by the clusters lies in one of the dendrogram’s subtrees. Due to how the dendrogram is constructed, a cell may end up being split off into its own cluster and is temporarily labeled as a non-classified “Singleton.” The merge step cleans up singletons and small clusters, resulting in a higher ARI. For the Yan et al.^9^ dataset, the ARI increases dramatically after the merge step. This is due to the fact that after the split step, 1) 15 of the 124 cells ended up as singletons, and 2) splitting generated twice as many clusters as needed. In fact, for this dataset, dividing each true cluster into two equal-sized parts would result in an ARI of 0.74 when compared with the original labels. A more detailed visual analysis of the Yan et al. dataset is given in Supplementary Figure 1. Under certain conditions, some cells may remain in their own clusters even after the merge step (see Implementation section). These cells are analogous to outliers.

**Figure 3.**
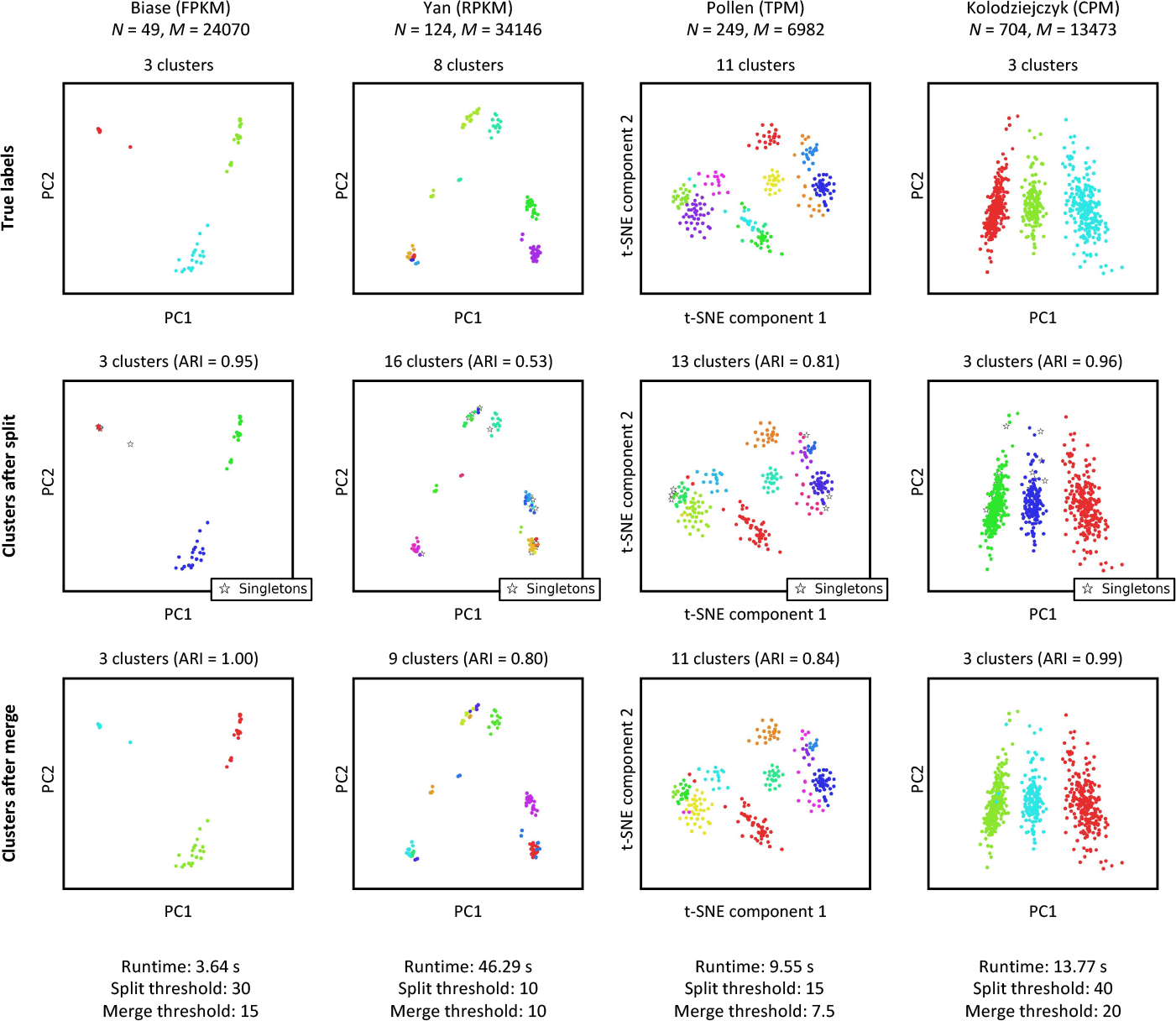
Gold standard datasets. DendroSplit is evaluated on four single-cell RNA-Seq datasets where the labels are highly likely to be correct^2,5,8,9,34^. In addition to visual inspection, cluster quality is evaluated using the adjusted Rand index (ARI) based on the true labels. We observe here that the split step tends to generate more clusters than expected, shrinking the ARI. Additionally, due to how the dendrogram is constructed, a cell may end up in its own cluster and is consequentially labeled as a “Singleton.” The merge step treats both these cases. The cells are visualized using either the first two principal components (PC) or the first two t-distributed stochastic neighbor embedding^63^ (t-SNE) components. The reported runtimes include computation of the pairwise distance matrices.

#### Exploratory analysis

We further demonstrate the exploratory power of DendroSplit on Patel et al.’s^7^ dataset of 430 cells, 5948 features from five primary human glioblastomas. Without any further preprocessing, DendroSplit recovers five clusters corresponding to the five glioblastomas (Figure 4A). Furthermore, DendroSplit can justify its findings by showing us the gene that plays the largest role in validating each split. Splits 4 and 5 in Figure 4B show distinctively how *SEC61G*, for example, distinguishes MGH26 cells from MGH29 and MGH31 cells. The analysis also gives insight on how the hierarchical clustering was performed. The cells from MGH26 were split in half during earlier stages of clustering, which is why they end up in separate superclusters at the root node. This is an artifact of the greedy nature of hierarchical clustering where clusters that should be close together may end up far apart. Merge 1 in Figure 4C shows DendroSplit fixing this. At the same time, we see that *PAN3* may be a valid marker for distinguishing these two subtypes within MGH26 cells. Further analysis and perhaps side information would be needed to decide whether or not these two subtypes are truly different. DendroSplit handles the subjectiveness associated with clustering by showing the factors that contribute to its decisions.

**Figure 4.**
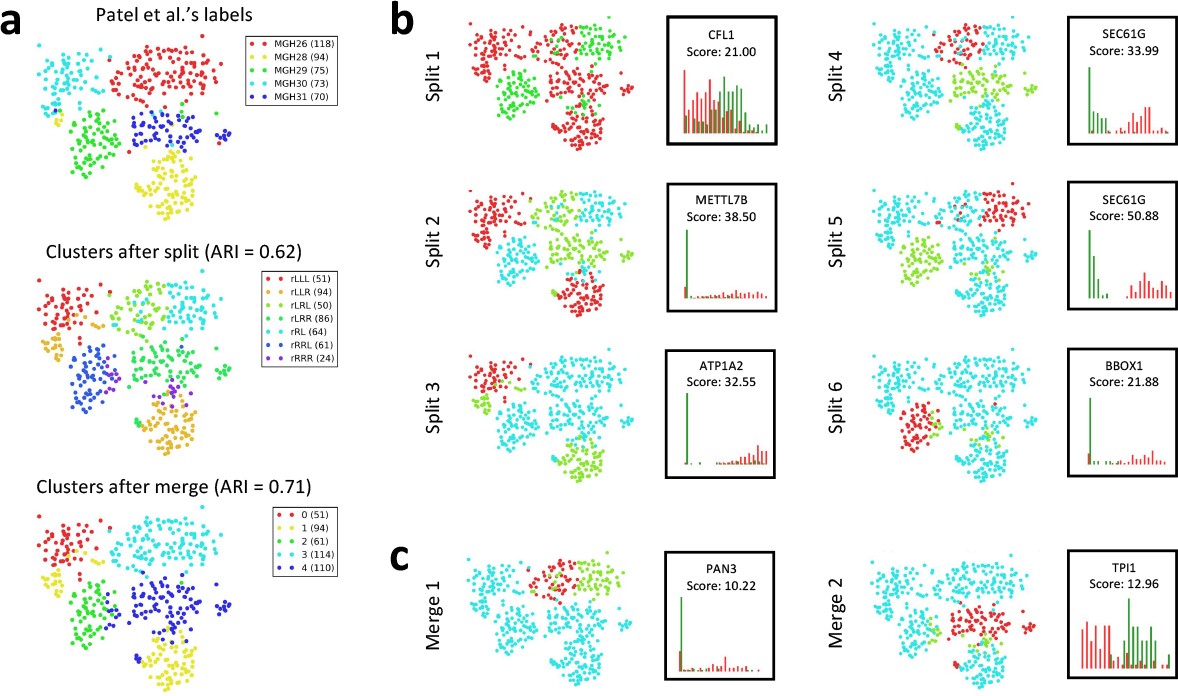
Exploratory analysis on Patel et al. dataset. **(a)** DendroSplit is evaluated on Patel et al.’s dataset of 430 cells, 5948 features (genes) from five primary human glioblastomas^7^. Gene expression is quantified using TPM. The split and merge thresholds are 20 and 15, respectively, and the analysis takes 9.64 seconds to run. The numbers in the legends represent the number of points in the corresponding clusters. For the split step, the names of the clusters are generated based on the position of the subtrees in the dendrogram, “r” represents the root node, and “rRL” represents the subtree found at the left child of the right child of the root, **(b)** We can evaluate how cells were partitioned at each step of the split procedure, and DendroSplit can also show us the within-cluster distributions of the gene that validates the split, **(c)** We can also evaluate how clusters obtained after the split step were combined during the merge procedure, and DendroSplit can show us the distributions of the most distinguishing gene between two merged clusters.

### Performance on larger single-cell datasets

We use DendroSplit to re-analyze three large single-cell RNA-Seq datasets that utilize unique molecular identifiers (UMIs) for quantifying genes^12, 17, 20^. Unlike for previous single-cell RNA-Seq datasets, the labels for these datasets were assigned using diverse computational methods. Figure 5 first shows that performing a feature selection step prior to analysis with DendroSplit decreases the runtime dramatically. In fact, for the Zeisel et al. dataset, filtering out genes using the procedure described by Macosko et al. reduces both *M* and the runtime by a factor of 16.6. Additionally, the filtering out of noisy features improves the quality of the distance metric, and we see that the ARI improves dramatically. We also report that using a much smaller split threshold of 15 results in 43 non-singleton clusters. When compared with Zeisel et al.’s 47 subclasses, we achieve an ARI of 0.42. The gene filtering procedure is used for the all datasets presented in Figures 5 and 6.

**Figure 5.**
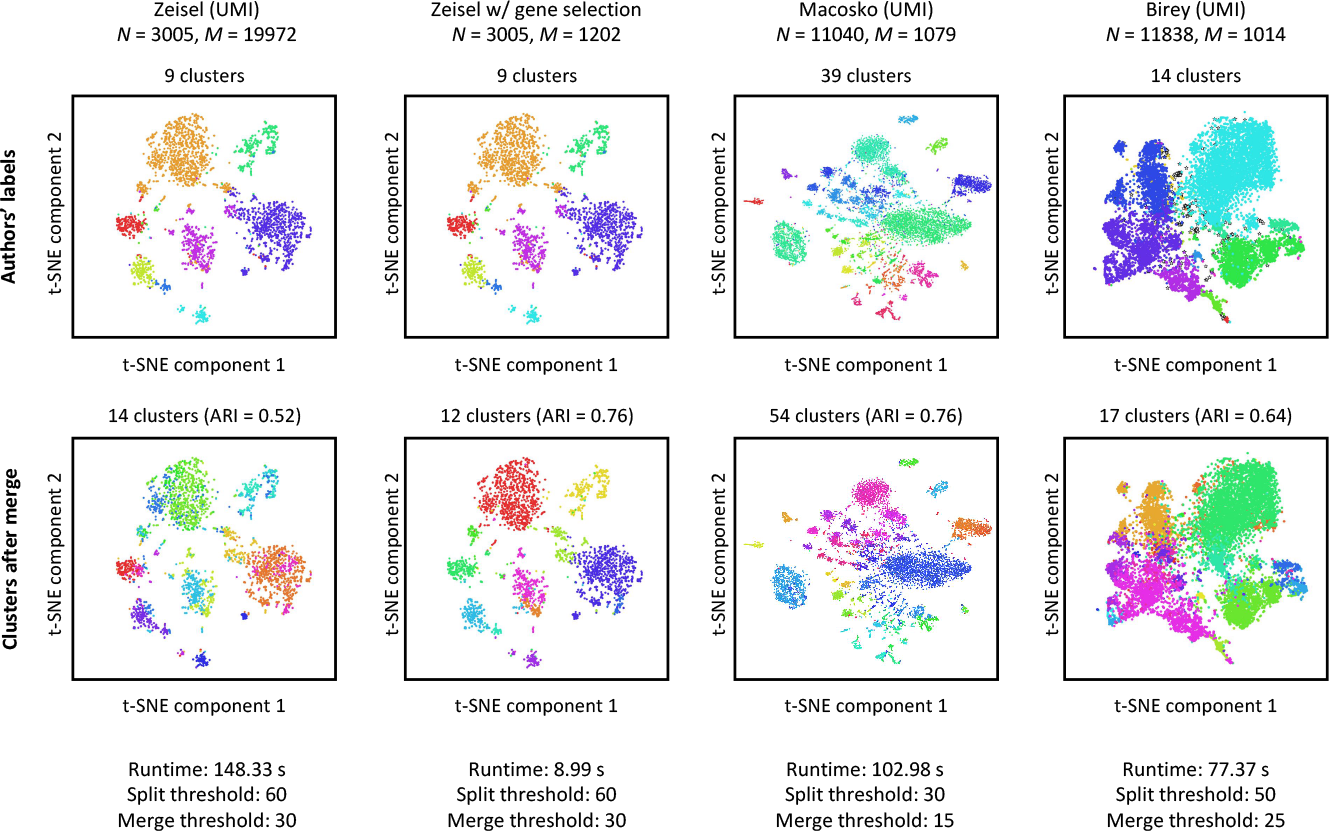
Larger datasets. DendroSplit is evaluated on three large single-cell datasets where the labels were assigned using computational methods^12,17,20^. Because *M* independent Welch’s i-tests are performed at each potential split, the runtime of the algorithm scales linearly with *M*. As demonstrated with the Zeisel et al. dataset, decreasing *M* by a factor of 16.6 likewise decreases the runtime by the same factor. The datasets here are preprocessed by filtering out genes using the procedure described by^20^. This preprocessing step also improves the quality of the distance metric used, resulting in better performance.

For the three datasets analyzed in Figure 5, DendroSplit generates similar but not identical labels. Figure 6A shows that DendroSplit disagrees even more strongly on Zheng et al.’s dataset of 17426 cells, 908 features from fresh peripheral blood mononuclear cells (PBMCs). Noting that the merge step does not increase the ARI significantly, we focus on the split step labels. Although 15 valid splits were recorded, we investigate only the 5 shown in Figure 6B. For the remaining splits, see Supplementary Figure 2. Split 1 was validated due to a lack of expression of several genes (*FCGR3A, LY86, FCN1*, and *IFI30*) in the red population, which we match to the authors’ CD34+ cells. Split 2 shows the separation of the red cells from the green cells based on high expression of *NKG7* and *GNLY*, markers for natural killer (NK) cells. The green cluster in split 5 likely corresponds to cytotoxic T cells based on increased expression of *GZMH*. The red cluster in split 9 shows greater expression of *CD79A* and may therefore represent B cells. Finally, the red cluster in split 14 does not have an obvious match with any of Zheng et al.’s set of labels. DendroSplit shows us that the existence of this cluster is justified based on increased expression of several genes including *FCGR3A, CFD*, and *LST1*. A one-versus-rest differential expression analysis based on independent Welch’s *t*-tests (see Supplementary Table 1) further shows that *PSAP* and *SERPINA1* are also overexpressed, indicating that the red cluster (cluster 6 after the merge step in Figure 6A) corresponds to some type of monocyte. We also repeat this analysis with the full 68579 cells, 20374 genes dataset, and the results are shown in Supplementary Figure 3.

**Figure 6.**
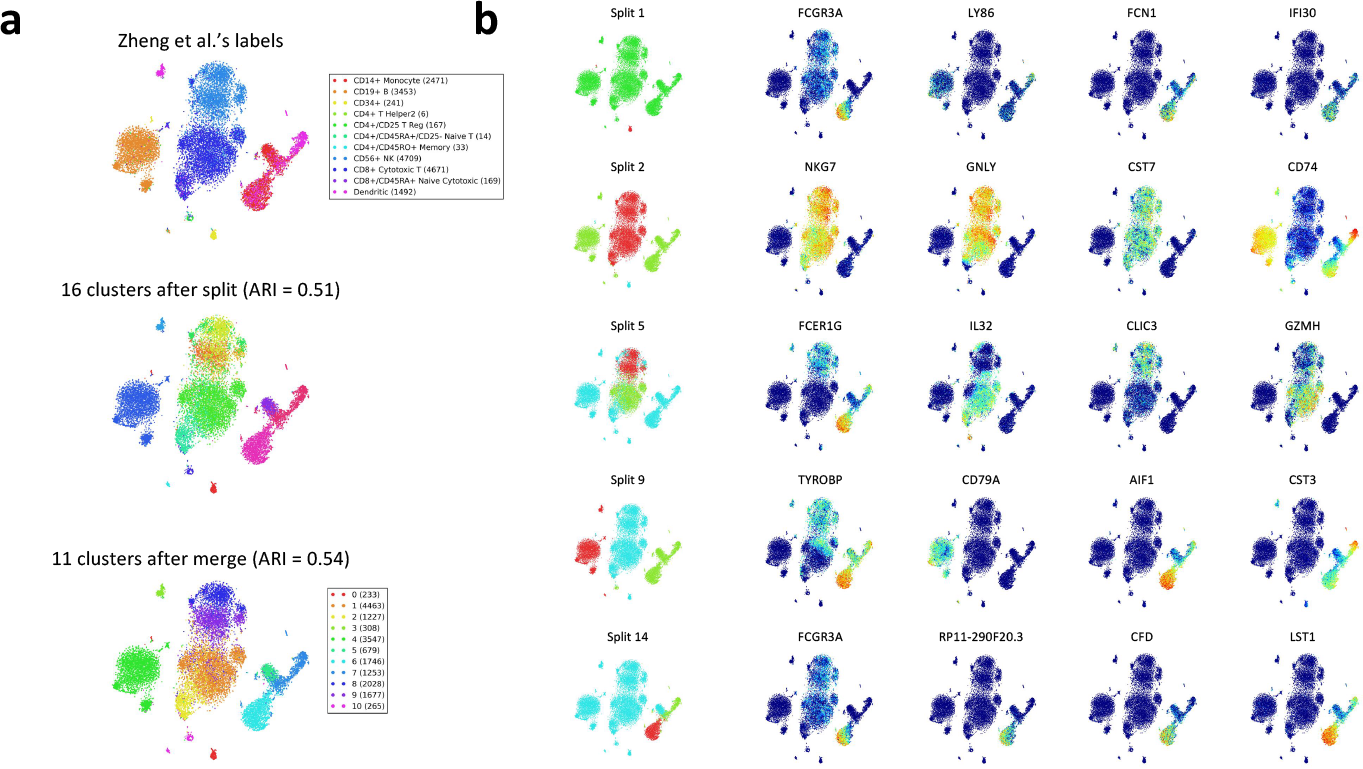
Exploratory analysis on PBMC dataset. **(a)** After gene selection and removal of cells with less than 50 counts across all genes, DendroSplit generates clusters for Zheng et al.’s remaining dataset of 17426 cells, 908 features (genes) from fresh peripheral blood mononuclear cells (PBMCs)^21^. Gene expression is quantified using UMI counts. The split and merge thresholds are 200 and 100, respectively, and the analysis takes 119.97 seconds to run. **(b)** 5 of the 15 recorded valid splits are shown along with the expression levels of the top 4 genes used for validating each split. The reported runtimes include computation of the pairwise distance matrices.

Finally, Figure 7 demonstrates the score threshold sweeping procedure for the Kolodziejcyk et al. and Zeisel et al. datasets. As observed in the experiments, larger datasets often require larger thresholds due to the *t*-statistic generally increasing with *N*.

**Figure 7.**
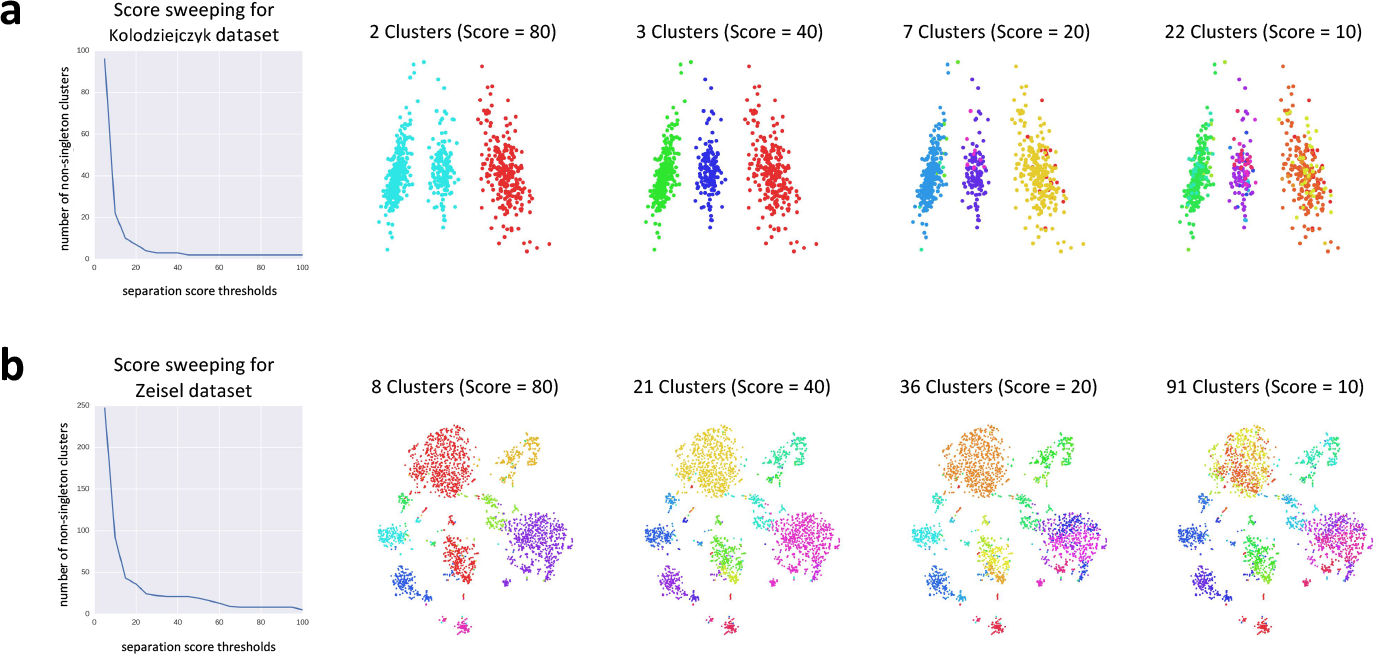
Split threshold sweeping. Since DendroSplit saves all relevant information at each valid split (e.g. *p*-values from Welch’s *t*-tests and IDs of cells being compared), we can run the method with a small split threshold to gather information about several potential splits. From this information, we can generate the labels we would have obtained after the split step had we run the algorithm with a larger split threshold. **(a)** For the 704 × 13473 Kolodziejczyk et al.^5^ dataset, running DendroSplit using a split threshold of 2 takes 37.63 seconds, and generating a set of new labels takes 0.032 seconds. **(b)** For the 3005 × 1202 Zeisel et al. dataset, running DendroSplit using a split threshold of 2 takes 13.48 seconds, and generating new labels takes 0.403 seconds. For both datasets, the DendroSplit runtimes include computation of the pairwise distance matrices.

## Conclusion

In this work, we presented a novel interpretable framework for tackling the single-cell RNA-Seq clustering problem. We demonstrated that a dendrosplit-splitting approach based on a separation score was key for uncovering the multiple layers of biological information within a dataset. In addition to recovering results from a diverse set of single-cell studies, we showed that the framework could cheaply produce several clusterings of the same dataset. Most importantly, the algorithm could justify each of its decisions in an interpretable way. Thus, DendroSplit is suitable as a backend algorithm for interactive analysis and interpretation.

With single-cell RNA-Seq technology improving, we can only expect increased cell throughput and larger datasets. While DendroSplit is able to generate clusters without expensive hyperparameter tuning, its optimal split and merge thresholds do depend on the size of the dataset since larger datasets tend to yield smaller *p*-values. To remove this size dependence, one could subsample a larger dataset to the same fixed size multiple times, run DendroSplit on each subsample, and ultimately report some consensus result. Another strategy for handling this dependence is in choosing a dataset-size-correcting statistical test rather than the naive Welch’s *t*-test when computing the separation score.

For the analyses in this work, we used a separation score based on a computationally cheap method of performing differential expression and a simple definition of cell type. Separation scores based on more complex methods of evaluating differential expression such as those presented by^31,45–51^ may yield better results at the cost of greater computation. Additionally, just like for other clustering approaches, existing tools including those designed for outlier detection^13, 52^, drop-out imputation^53^, and correcting other sources of technical noise^54–62^ can be easily incorporated into the DendroSplit framework by applying the desired correction procedures before the clustering step.

## Availability

**Project name**: DendroSplit

**Project home page**: https://github.com/jessemzhang/dendrosplit

**Operating system(s)**: Platform independent

**Programming language**: Python 2.7

**Other requirements**: Python modules `numpy 1.12.1, scipy 0.19.0, matplotlib 1.5.3, sklearn 0.18.1, networkx 1.11, community`

**License**: Creative Commons Attribution-NonCommercial-ShareAlike 4.0 International license

**Any restrictions to use by non-academics**: License needed

## Abbreviations

**ARI**: Adjusted Rand index

**backSPIN**: Back sorting points into neighborhoods

**DBSCAN**: Density-based spatial clustering of applications with noise

**NK**: Natural killer cells

**PBMC**: Peripheral blood mononuclear cells

**RNA-Seq**: Ribonucleic acid sequencing

## Declarations

### Ethics approval and consent to participate

Not applicable.

### Consent for publication

Not applicable.

### Availability of data and material

The^2,5,7‘9,12,17,20,21^ datasets were obtained from public repositories. The^5,8^ datasets were obtained from the GitHub repository for^30^.

### Author’s contributions

JMZ conceived the idea of splitting dendrograms and merging clusters using *p*-values, wrote the software package, performed analyses of data, interpreted results and wrote the manuscript. JF, HCF, DR, and DNT interpreted results, supervised the project and wrote the manuscript.

## Acknowledgements

We thank Govinda Kamath and Vasilis Ntranos for feedback and useful discussions about analyzing single-cell RNA-Seq datasets.

## Additional files

1. `bmc_bioinformatics supplement.pdf`. **Supplementary figures and separation score analysis**. A file containing a distance-metric interpretation of separability score along with the three supplementary figures described in the main text: 1) visual analysis of the Yan et al. dataset, 2) further analysis for the splits generating for the Zheng et al. dataset, 3) analysis on the full Zheng et al. dataset.
2. `Supplementary Table 1.tsv`. **PBMC differential expression analysis**. A one-versus-rest differential expression analysis based on independent Welch?s t-tests for the Zheng et al. PBMC dataset of 17426 cells, 908 features described in the main text.

